# Genotype imputation using the Positional Burrows Wheeler Transform

**DOI:** 10.1101/797944

**Authors:** Simone Rubinacci, Olivier Delaneau, Jonathan Marchini

## Abstract

Genotype imputation is the process of predicting unobserved genotypes in a sample of individuals using a reference panel of haplotypes. In the last 10 years reference panels have increased in size by more than 100 fold. Increasing reference panel size improves accuracy of markers with low minor allele frequencies but poses ever increasing computational challenges for imputation methods.

Here we present IMPUTE5, a genotype imputation method that can scale to reference panels with millions of samples. This method continues to refine the observation made in the IMPUTE2 method, that accuracy is optimized via use of a custom subset of haplotypes when imputing each individual. It achieves fast, accurate, and memory-efficient imputation by selecting haplotypes using the Positional Burrows Wheeler Transform (PBWT). By using the PBWT data structure at genotyped markers, IMPUTE5 identifies locally best matching haplotypes and long identical by state segments. The method then uses the selected haplotypes as conditioning states within the IMPUTE model.

Using the HRC reference panel, which has ~65,000 haplotypes, we show that IMPUTE5 is up to 30x faster than MINIMAC4 and up to 3x faster than BEAGLE5.1, and uses less memory than both these methods. Using simulated reference panels we show that IMPUTE5 scales sub-linearly with reference panel size. For example, keeping the number of imputed markers constant, increasing the reference panel size from 10,000 to 1 million haplotypes requires less than twice the computation time. As the reference panel increases in size IMPUTE5 is able to utilize a smaller number of reference haplotypes, thus reducing computational cost.

**Author summary:** Genome-wide association studies (GWAS) typically use microarray technology to measure genotypes at several hundred thousand positions in the genome. However reference panels of genetic variation consist of haplotype data at >100 fold more positions in the genome. Genotype imputation makes genotype predictions at all the reference panel sites using the GWAS data. Reference panels are continuing to grow in size and this improves accuracy of the predictions, however methods need to be able to scale to increased size. We have developed a new version of the popular IMPUTE software than can handle referenece panels with millions of haplotypes, and has better performance than other published approaches. A notable property of the new method is that it scales sub-linearly with reference panel size. Keeping the number of imputed markers constant, a 100 fold increase in reference panel size requires less than twice the computation time.

## Introduction

Genotype imputation is a widely used method in human genetic studies that infers unobserved genotypes in a sample of individuals. In a typical scenario, the study samples are genotyped on a SNP microarray with between 300,000 to 5 million markers. This data is then combined with a reference panel of haplotypes with many tens of millions of markers, and a statistical model is used to predict the genotypes at these markers in the study samples [1].

One of the main applications of genotype imputation is to increase the resolution of genome-wide association studies (GWAS). Imputed datasets increase the number of markers that can be tested for association. For example, in the UK Biobank dataset [2] imputation increased the number of testable markers from 825,927 to over 96 million. This increased number of SNPs can boost the power of the study. Genotype imputation also facilitates meta-analysis across cohorts that are often genotyped using different SNP microarrays. Imputatation on from the same reference panel standardizes the set of testable markers, allows simple integratation of data and/or results across studies [3]. Imputation can also be used to predict markers necessary to calculate polygenic risks scores (PRSs), which typically involve a weighted sum of genotypes across the genome.

Many different methods have been proposed over the years [4], however the most widely used and accurate methods are based on Hidden Markov Models (HMM). Typically the study samples will have been phased in advance using accurate methods [5–7], and this has become known as ‘pre-phasing’ [8]. The imputation HMM is then used to model the sharing of sequence between the haplotypes in the study sample (which we refer to as the *target haplotypes*) and the haplotypes in the reference panel. The HMM models each study haplotype as an imperfect mosaic of haplotypes in the reference panel [1]. The output of the HMM at each position in the genome is a vector of copying probabilities for each haplotype in the reference panel. For each of the markers in the reference panel these are used to make a weighted prediction of the unobserved allele at the markers.

One of the most important factors that determines imputation quality is the number of haplotypes in the reference panel. As the number of haplotypes increases, each study sample haplotype is able to find fewer longer stretches of matching sequence in the reference panel, which increases the accuracy. Table 1 shows how reference panel size has increased over the years due to projects such as the International HapMap Project [9], the 1000 Genomes Project (1000G) [10], the UK10K Project [11], and the Haplotype Reference Consortium (HRC) [12]. Soon larger reference panels will become available from the Trans-Omics for Precision Medicine (TOPMed) program [13] and the 100,000 Genomes Project [14], both of which will exceed 100,000 high-coverage whole genome sequenced samples. Further ahead, as sequencing data on all 500,000 participants of the UK Biobank [2] becomes available, this will be used as an even larger reference panel.

**Table 1.** Evolution of commonly used imputation reference panels over time.

| Reference panel | Released Year | Number of Samples | Number of Markers (Millions) |
| --- | --- | --- | --- |
| HapMap Project phase 3 | 2009 | 1,011 | 1.4 |
| 1000 Genomes phase 1 | 2012 | 1,092 | 29.0 |
| 1000 Genomes phase 3 | 2014 | 2,504 | 81.7 |
| UK10K Project | 2015 | 3,781 | 42.0 |
| HRC | 2016 | 32,470 | 40.4 |

In this paper we present IMPUTE5, a genotype imputation method designed to handle the new generation of reference panels. To achieve this, the method builds on three main components: (i) the use of a new reference panel file format, allowing fast access to data in specific chromosome regions (ii) the use of the PBWT [15] to select a subset of reference panel haplotypes and reduce the state space in the IMPUTE model [16] (iii) imputation during output directly into the BGEN [17] file format that is specifically designed for imputation data. To demonstrate the superior performance of IMPUTE5, we benchmark our imputation method against IMPUTE4 [2], MINIMAC4 [18] and BEAGLE5.1 [19], using simulated reference panels up to 1,000,000 haplotypes in size, and real reference panels such as the 1000 Genomes project reference panel [10] and the Haplotype Reference Consortium [12].

## Materials and Methods

### PBWT

The main methodological advance in this paper is the incorporation of the PBWT [15] into the IMPUTE model. This section provides some brief background on the PBWT needed to describe the IMPUTE5 method. The PBWT is a generic way to encode binary matrices, especially useful in the case of haplotypes at a set of binary markers, each with two alleles arbitrarily coded as 0 and 1. Let *H* = {*h*_0_*, h*_1_, … , *h*_*N*−1_} be a set of *N* haplotypes genotyped at *M* markers, *h_n_* = {*h*_*n*,0_, *h*_*n*,1_, … , *h*_*n,M*−1_}, where *h*_*n,m*_ ∈ {0, 1}, represents the *n*th haplotype. *H* can be thought as a *N* × *M* binary matrix: each entry of the matrix is defined by the reference haplotype (row) and marker (column). For this reason, we refer to *H* using the usual matrix notation.

The PBWT of *H*, indicated as *Y*, is another *N* × *M* binary matrix, where the *m*-th column of *Y* is an invertible transformation of the *m*-th column of *H*. In its basic form, *Y* is complemented by another *N* × (*M* + 1) matrix *A*, where every column *A*_:,*m*_, called the *positional prefix array*, is a permutation of {0, … , *N* − 1} which defines the reverse prefix order of the haplotypes in *H* up to marker *m* − 1.

Using a similar notation as in [20], we define the binary string 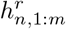 as the reverse prefix of the *n*-th haplotype ending at marker *m*:

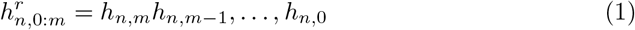

and let be 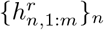 be the set of all the *N* reverse prefixes at marker *m*. We then define *A_n,m_* to be the index of the *n*-th lexicographically sorted reverse prefix along the set 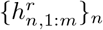. *A*_:,*m*_ represents a bijection on {0, … , *N* − 1} and thus is invertible. As a special case, we define *A_n,−_*_1_ = *n*, representing the order of empty reverse prefixes.

The PBWT of *H* is directly derivable from *H* and the prefix array *A*:

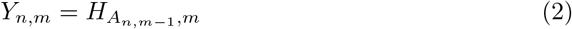

in other words, the PBWT at marker *m* is the vector of values of the haplotypes in *H* at marker *m*, (*H*_:,*m*_), in the order defined by the reverse prefix array at marker *m* − 1, *A*_:,*m−*1_.

One use of *Y*_:,*m*_ is to update *A*_:,*m*−1_ to *A*_:,*m*_. Suppose *b* is a symbol, *b* ∈ {0, 1}. We can define a mapping between positional prefix array at markers *m* − 1 and *m*:

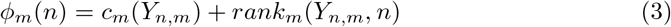

where *c_m_*(*b*) gives the number of symbols in *Y*_:,*m*_ that are lexicographically smaller than *b* and *rank_m_*(*b, n*) the number of *b* symbols in *Y*_:,*m*_ before position *n*. The *n*-th haplotype in the positional prefix order at column *m* − 1 is ranked *φ_m_*(*n*) in column *m*. Thus

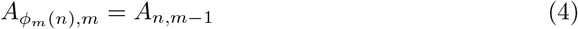

Eqs (2 and 4) give a procedural algorithm to compute *A*_:,*m*_ and *Y*_:,*m*_ from *H*_:,*m*_ and *A*_:,*m*−1_. Since there is strong correlation between adjacent markers in *H* due to linkage disequilibrium, there are long runs of the same symbol in the columns of *Y*. This makes columns of *Y* much more compressible than the columns of *H*.

*Y* and *A* represent only the basic form of the PBWT. It is possible to complement them by storing additional information such as the rank indices *U, V* (usually called FM index [21]). Each column *m* of these two matrices store information about the *rank_m_*(*b, n*) for symbol *b* = 0 and *b* = 1 respectively. It is also important to notice that there is no need to store both these matrices because it is possible to derive one from the other: *rank_m_*(1 − *b, n*) = *n* + 1 − *rank_m_*(*b, n*) since *rank_m_*(0, *n*) + *rank_m_*(1, *n*) = *n* + 1.

Another important information that can be added is the divergence matrix *D*. Columns of *D* contain the position of the last (reverse prefix) mismatch between adjacent haplotypes in the order *A*. The value of *D_n,m_* is defined to be the smallest value *m*′ such that 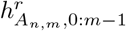 matches 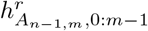. In the case of a mismatch, the value of *D*_*n,m*_ is set to *m*. An important property is that the start of any maximal match ending at *m* between any 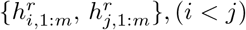 is given by:

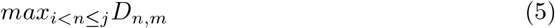

The cost of building a PBWT *Y* from *H* is *O*(*NM*), including all the complementary matrices described above. Using the PBWT indices, it is possible to find maximal matchings within *H* in linear time and find maximal matchings of a new sequence *z* in *H* in *O*(*M*), independently from the number of haplotypes in *H*.

### IMPUTE model

IMPUTE5 is a haploid imputation method, assumes that both the reference and study samples are phased and contain no missing alleles at any site. In what follows we will refer to the phased study samples as the *target panel* of haplotypes. IMPUTE5 uses the same HMM used in previous versions of the IMPUTE software [16] that is based on the Li and Stephens model [22]. Each reference haplotype represents a hidden state of the HMM. The model assumes that each target haplotype is an imperfect mosaic of haplotypes emitted from the sequence of hidden states representing the reference panel haplotypes. The changes from one state to another are modelled as recombination events and the observed target allele may differ from the alleles on the underlying true haplotypes to allow for mutation and genotype error.

#### HMM definition

Let *H* be the set of *N* haplotypes genotyped at *M* markers, that have been selected as a subset from a reference panel of haplotypes. The way in which the *N* haplotypes are chosen in each window is described in a later section. We also have a set of *K* study sample (target) haplotypes, defined only at a subset of the *M* markers. We refer to the set of *T* markers that are genotyped in both the panels as *target markers* 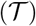, and the others, present only in the reference panel, as *reference markers* 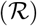. Consecutive pairs of haplotypes represent the diplotype of each study individual. We define the HMM model only at target markers. Therefore, we use the symbol 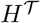 to indicate the restriction of the reference panel *H* to target markers and we use the symbol *m* to indicate a marker in 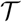.

Given a target haplotype *t* the probability of observing *t* from 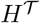 can be then written as:

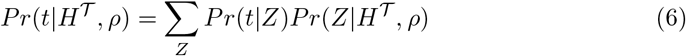

where *Z* is a sequence of unobserved copying labels, one for each target marker, *Z_m_* ∈ {0, 1, … , *N* − 1} and the term 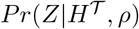 models sequence of transitions of the HMM and is defined by

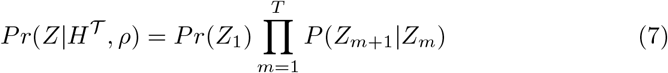

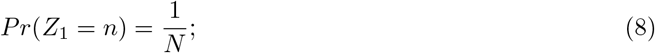

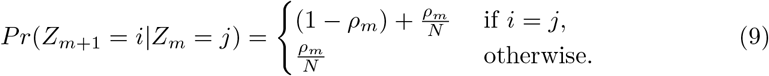

where *ρ_m_* is a locus specific parameter modelling genetic recombination events, defined as 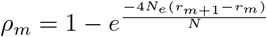, where *N_e_* is the effective diploid population size and *r*_*m*+1_ − *r_m_* is the average rate of crossover per unit physical distance per meiosis between target markers *m* + 1 and *m* multiplied by their physical distance. Eq (9) is motivated by the fact that recombination events can be described as a Poisson process having rate 4*N_e_*(*r*_*m*+1_ − *r_m_*)/*N*.

We model the emission probability *Pr*(*t*|*Z*) in Eq (6) differently to the standard IMPUTE model, and we adopt a simpler version as in the SHAPEIT4 model [6], reducing the equation to:

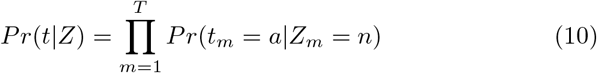

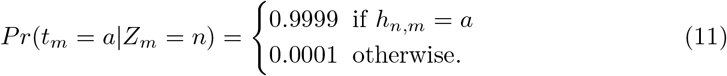

where *a* ∈ {0, 1} is a haplotype value. It has been shown that imputation is relatively insensible to the mutation parameter [23], and we tested that the new emission probability slightly increases accuracy, especially in the case of big reference panels.

#### Imputation

The posterior probability of the hidden states is computed using the forward-backward algorithm [24]. IMPUTE5 calculates and stores these quantities at each target marker. The imputation step is performed after the marginal posterior distribution of the copying states has been computed. The state probabilities at reference markers 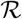 can be linearly interpolated from the probability at the two bounding target markers. The motivation of using linear interpolation is, that over short genetic distance, the change in state probabilities can be approximated by a straight line [2, 23]. The imputed probability for a particular allele is then just the sum of all the state probabilities at that marker in which the correspondent reference haplotypes carry the allele.

As performed by BEAGLE5 [19], we store state probabilities at consecutive markers for a reference haplotype only if one of the state probabilities is greater than the inverse of the number of states considered in the HMM. Since only a small subset of the state probabilities at consecutive markers needs to be stored, imputation can be delayed during output, saving the memory required to store imputed probabilities at reference markers.

At the end of the forward-backward pass, a small subset of posterior probabilities are stored at each target site. When performing imputation, IMPUTE5 exploits the fact that, when imputing from large reference panels, a sizeable fraction of the imputed variants will be imputed as monomorphic and therefore imputation could be avoided for these markers. If the variant is rare, a simple test is performed to verify that at least one of the thresholded states carries the alternative allele in the reference panel. If that is the case, standard imputation is performed at the marker, otherwise a monomorphic variant is printed in output and no additional computation is required. We refer to this as delayed lazy imputation. This simple procedure has an impact in the case of big reference panels containing a large number of rare variants.

In addition to this, IMPUTE5 does not store the reference panel at imputed variants in memory, but they are streamed by reading the reference panel during imputation. This, combined with delayed lazy imputation, allows quick imputation from extremely large reference panels, consuming only a small amount of the memory.

### State Selection Using the PBWT

IMPUTE5 uses the PBWT of the reference panel at target markers to identify a subset of states that share long identity by state (IBS) sequences with target haplotypes. Using just a subset of haplotypes saves computation time memory usage. The copying state selection is performed upfront before the HMM calculations and determines the set of *N* haplotypes in 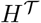. This set will be different for each target haplotype.

The PBWT of the reference panel at the target markers is calculated sequentially from left to right across the region being imputed and the state selection occurs at the same time. So after one pass through the full dataset the state selection has been performed for *all* the target haplotypes. This means that there is no need to store the full PBWT of the reference panel in memory.

The selection procedure occurs in two steps. First, each target haplotpes is inserted (or located) in the PBWT. Second, haplotypes ‘close’ to the target haplotype in the PBWT are identified. This selection only occurs at a relatively sparse set of target markers. Moving left to right through the PBWT the set of ‘close’ haplotypes are added to a list and this list is then used as the copying set of states in the HMM. Fig 1 illustrates the method on a small example dataset.

**Fig 1.**
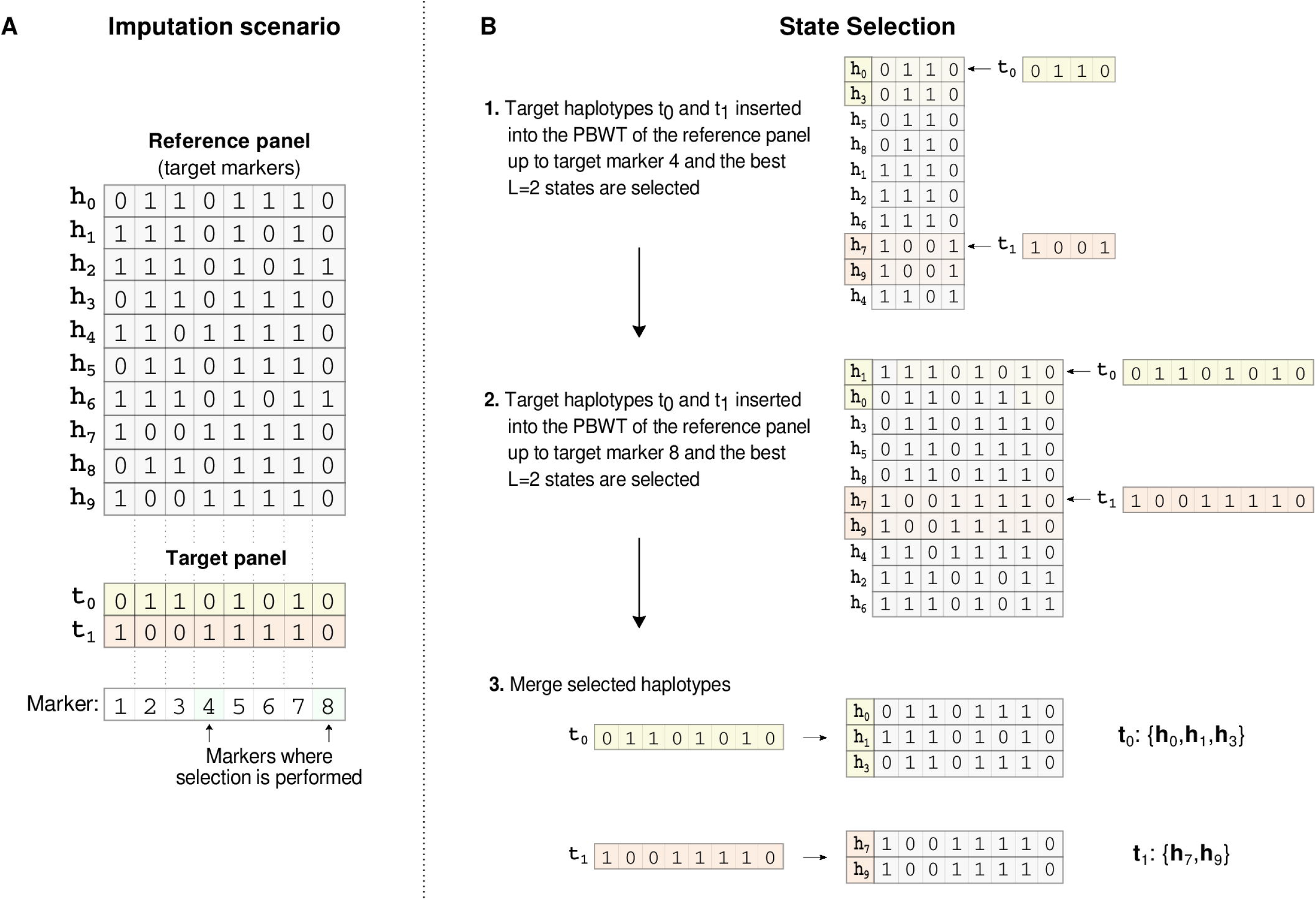
IMPUTE5 copying state selection Small example to illustrate IMPUTE5 copying state selection. (A) A reference panel of haplotypes *H* = {*h*_0_, … , *h*_9_} is restricted to the set of target markers and is shown together with a target panel of two haplotypes *T* = {*t*_0_, *t*_1_}. The copying state selection is only performed at a subsetof target markers. In this example, these are the 4th and 8th markers, and are shaded green. (B) The target haplotypes are inserted into the PBWT, using the rank operations (FM-index). In (B1) target haplotypes {*t*_0_, *t*_1_} are searched in the positional prefix array of the reference panel up to marker 4 and *L* = 2 reference haplotypes are selected for each target haplotype. In (B2) *t*_0_ and *t*_1_ are searched in the positional prefix array of the reference panel up to marker 8 and again *L* reference haplotypes are selected. (B3) The selected haplotypes are then merged to form a list of copying states for each target haploype. The list may not necessarily be the same length. These states will be used in the HMM to perform imputation.

Inserting each target haplotype into the PBWT involves searching the prefix array of the reference panel using Eq (3) and the FM-index. For a target *t* at marker *m*, this search finds the location of the reference haplotype in the PBWT that shares the longest reverse prefix with *t* up to marker *m*.

The updated matched position *f* of the target *t* at marker *m* is given by:

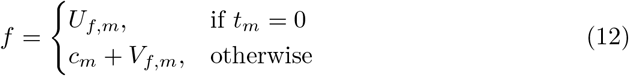

where *c_m_*, *U*_:,*m*_ and *V*_:,*m*_ are respectively the number of 0s in each marker *m* and the rank matrices for the reference panel at marker *m*. The search itself costs *O*(1) at each marker using the FM-index of the PBWT and it is therefore independent of the number of haplotypes in the reference panel.

We keep the list of locations for each target haplotype. At each marker, we first update the PBWT of the reference marker and then we update the list of target locations. The selection is performed every interval of length *I* (0.02 cM by default). We call *selection markers* the markers where the selection is performed. The cost of the search is *O* ((*N* + *K*)*M*), where *K* is the number of target haplotypes.

#### Selection algorithms

IMPUTE5 has two algorithms that select the ‘closest’ *L* haplotypes to the target haplotype within the PBWT. Both selection methods have *O*(*LK*) computational cost. The first one, which we call *divergence selection*, and first proposed in the software package SHAPEIT4 [6], selects the best *L* states in the neighbourhood of the current match position *f* by using the divergence matrix, exploiting Eq (5).

By design the PBWT encapsulates a large amount of local linkage disequilibrium information. The longest reverse prefixes are by definition in the neighbourhood of the best matching haplotype found during the search. In order to take only the best haplotypes, the divergence matrix is used.

Starting from an optimal position *f* at marker *m*, it checks the values of the divergence array at *i* = *f* − 1 and *j* = *f* + 1. If *D_i,m_* <= *D_j,m_* then *i* is decreased and positional prefix *A_i,m_* is added to the list of selected states, otherwise *j* is increased and *A_j,m_* is added to the list. The algorithm continues until *L* states are selected. A pseudo algorithm of the copying state selection is shown in S1 Algorithm.

The second selection algorithm, which we call *neighbour selection*, does not use the divergence matrix. At every selection marker, it simply takes the *L* neighbouring states (*L/*2 in both the directions) of the current best match position. This algorithm only guarantees to select the best *L/*2 reverse prefixes, but it requires less operations than the divergence selection, because it does not need to compute and interrogate the divergence array for that marker. The only checks needed are when the target haplotype occurs close to a border (start or end of the positional prefix array), and to avoid copying a mismatch position. In the case that the target haplotype is close to a border, less than *L* states are selected for that marker. A pseudo algorithm of the copying state selection is shown in S2 Algorithm.

Durbin et al. [15] proposed an algorithm to find the set of set maximal matches with the target haplotypes. The set of set maximal matching is the set of states that share the longest stretches with the target haplotype. We tested the use of the set maximal matches as a selection algorithm and we found that these contain a lot, but not all of the relevant information, and there is a small but evident loss in accuracy when we use only those matches.

#### IMP5 File Format

We developed a new file format, called imp5 to read the reference panel quickly into memory. Each marker is stored independently in one of two different ways: if the alternative allele is rare (MAF < 1/256), the indices of the haplotypes that carry the alternative allele are stored, otherwise the sequence of alleles is stored using one bit per allele. Imp5 files are compact in memory and do not require other compression algorithms like gzip. This makes reading from a file an efficient operation, similar to bref3 [19].

The binary stored data structure coded in the imp5 file format is also used internally within IMPUTE5 to store the reference panel in memory. When imputing each target haplotype, at each target marker, the set of selected reference haplotypes that carry the alternate allele are needed. If the target site is stored as a bitset in the reference panel then the lookup is straightforward. If the site is stored as a list of indices of alternate alleles then either the list of reference panel indices is searched for the selected state index, or vice versa, depending upon which search is likely to be quicker.

Another feature of the imp5 files is that they are indexed, so that regions can be extracted efficiently. The indexing was developed along the same lines as *bgenix* [17], using sqlite3. The indexing is an important feature for imputation, especially when imputing different windows on the same chromosome independently. Other file formats like bref3 [19] and m3vcf [18] do not provide an index so far and therefore cannot directly interrogate arbitrary regions in constant time. In addition to this, IMPUTE5 can also read reference panels stored in VCF/BCF format.

IMPUTE5 requires that the reference and the target panel files are indexed, using the native imp5 index or tabix in the case of VCF/BCF files. In this way, several independent imputation jobs can be run at the same time, using a multi-process parallelization approach. A comparison of the memory required to store m3vcf, bref3 and imp5 file formats is given in Table 2.

**Table 2.** Memory (GB) required by reference file formats.

| Reference panel | vcf.gz | bcf | m3vcf.gz | bref v3 | imp5 |
| --- | --- | --- | --- | --- | --- |
| 1000 Genomes chr 20 | 0.33 | 0.28 | 0.09 | 0.19 | 0.38 |
| 1000 Genomes chr 10 | 0.73 | 0.62 | 0.21 | 0.42 | 0.85 |
| HRC chr 20 | 1.70 | 1.50 | 0.29 | 0.61 | 1.90 |
| HRC chr 10 | 3.90 | 3.40 | 0.60 | 1.30 | 4.30 |
| Sim10K | 0.06 | 0.04 | 0.02 | 0.02 | 0.04 |
| Sim100K | 0.83 | 0.56 | 0.15 | 0.11 | 0.42 |
| Sim1M | 18 | 10 | 1.90 | 0.75 | 4.06 |

Memory usage in Gigabytes required to store the HRC and 1000 Genomes project reference panels (chromosome 10 and 20) and 10 Mb of reference sample data for 10K, 100K, and 1M simulated UK European reference samples. The reported memory value is the amount of space used to store the data in *vcf.gz*, *bcf*, *m3vcf.gz*, *bref3* and *imp5* file formats. For the *imp5* file format, the value reported is the sum of the memory required by the *imp5* file plus the index file. Imp5 has been optimised to provide random access and fast reading time for a region of the chromosome and not for data compression.

### Parallelization

A typical IMPUTE5 job runs on multiple 10-20cM regions in parallel. Each region is completely independent from the others and can be run on different machines. The use of the indexing of the IMP5 files allows each process to read the reference panel efficiently.

Output is written in VCF, BCF or in BGEN v1.2 file format [17], the latter explicitly designed to store imputed data. Concatenating VCF, BCF and BGEN files at the end of imputation is an efficient process and allows to impute each window independently (*bcftools concat* or *cat-bgen* commands are used to merge output files).

IMPUTE5 can also multi-thread each process. We developed multi-threading using a shared memory approach. Each thread is responsible for a single target haplotype when running the HMM, or an imputation region between two target markers. The data sharing approach is crucial for reducing the memory required by each computational thread.

### Real and simulated data experiments

We compared IMPUTE5 to other existing imputation methods using real reference panels from the 1000 Genomes Project [10] and the Haplotype Reference Consortium [12]. We used data from both chromosome 10 and chromosome 20. For both panels we extracted a subset of samples, thinned down to a subset of sites, that are uses as the target haplotype panel, and used the remaining samples as reference panel. We also use a UK-European reference panel of simulated data for 10K, 100K, and 1M samples generated using MSPRIME [25]. Details of the real and simulated datasets are summarised in Table 3.

**Table 3.**
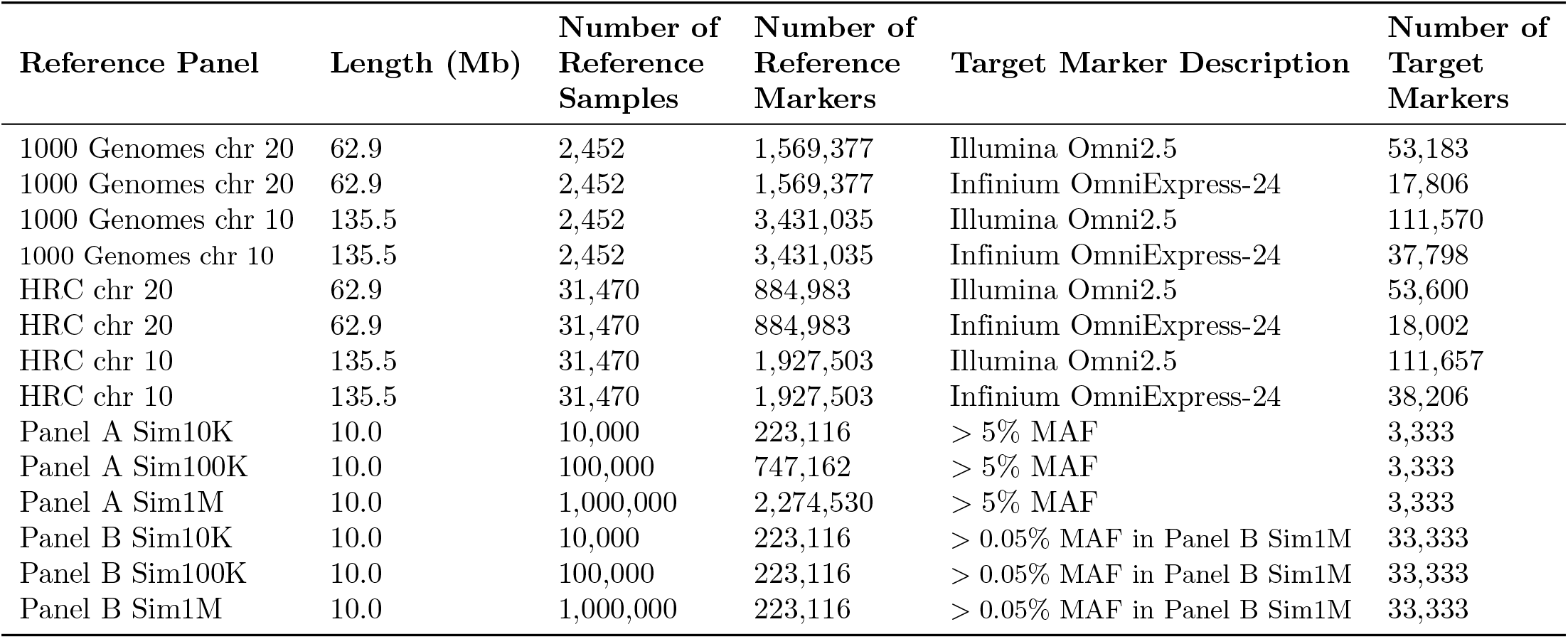
Summary of the real and simulated datasets used in comparing methods.

#### 1000 Genomes Project

The 1000 Genomes Project phase 3 dataset contains phased sequenced data of 2,504 individuals sampled from 26 different populations. As performed in the BEAGLE5 paper [19], we selected two random individuals from each population for the imputation target and used the remaining data as a reference panel. We restricted the 1000 Genomes reference data to markers having at least one copy of the minor allele in the reference panel, getting 3,431,035 markers on chromosome 10 and 1,569,377 markers on chromosome 20. In the 52 target samples, we masked markers that were not on the Illumina Omni2.5 array and the less dense Infinium OmniExpress-24 v1.2, resulting in 111,570 (Omni 2.5) or 37,798 (Infinium OmniExpress-24) target markers on chromosome 10 and 53,183 (Omni 2.5) or 17,806 (Infinium OmniExpress-24) target markers on chromosome 20. The list of markers has been obtained from https://www.well.ox.ac.uk/~wrayner/strand/.

#### The Haplotype Reference Consortium

Haplotype Reference Consortium (HRC) [12] reference panel combines sequence data across 32,470 individuals from 20 sequencing studies. We randomly selected 1,000 target individuals from the HRC panel and used the other 31,470 as a reference panel.

We removed monomorphic markers in the reference samples. In the target samples, we masked markers that were not on the Omni2.5 array and Infinium OmniExpress-24 v1.2, resulting in 111,657 (Omni2.5) or 38,206 (Infinium OmniExpress-24) target markers on chromosome 10 and 53,600 (Omni 2.5) or 18,002 (Infinium OmniExpress-24) target markers on chromosome 20.

In order to verify the sub-linear properties of IMPUTE5, we also randomly downsampled the HRC dataset to a subsets containing 30,000, 20,000, 10,000, 5,000, 3,000, 2,000 and 1,000 samples.

#### Simulated Reference Panels

We used MSPRIME [25] to simulate a 10Mb region of sequence data of UK-European samples. We simulated 11,000, 101,000 and 1,001,000 samples and extracted 1,000 samples from each of the three dataset, in order to have three reference panels of size 10K, 100K and 1M samples. We split each of the 1,000 target samples into three different target panels of size 10, 100 and 1,000.

In the target panels, we masked all but 3,333 markers, randomly selected between the markers having MAF > 5%, to simulate chip sites. The reference panels have 223,116, 747,162 and 2,271,530 markers respectively. We refer to this setting (reference panel + target panels) as Panel A.

We created 3 other simulated datasets (called Panel B), with 1 million, 100,000 and 10,000 samples, and each with the same number of 223,116 markers. We created 3 target panels of size 10, 100 and 1,000 samples at a subset of 33,333 markers by randomly selecting markers having MAF > 0, 05% in the 1M reference panel. Panel B is used to benchmark imputation on the same set of markers, varying the size of the reference panels.

## Results

### Comparison of Methods

We compared IMPUTE5 to IMPUTE4, MINIMAC4 (v.1.0.0) [18] and BEAGLE5.1 (version 25Nov19.28d) [19]. For simulated datasets, we used default parameters for each program. For real datasets used different imputation window size, depending on the imputation program. For IMPUTE4 we used imputation regions of 5Mb and 500kb of buffer, as perfomed for the UK Biobank imputation [2]. For MINIMAC4 we used the default settings (20Mb region). We run IMPUTE5 and BEAGLE5.1 on the same regions of 20 cM. In this case we used a 1Mb buffer region for IMPUTE5 and 2cM buffer region for BEAGLE5.1.

We used the HapMap2 [9] genetic map for BEAGLE5.1 and IMPUTE5 for real data imputation and the true genetic map for analyses with simulated data. MINIMAC does not require a genetic map, as recombination parameters are estimated and stored when producing the m3vcf format input file for the reference data.

BEAGLE5.1, MINIMAC4 and IMPUTE5 use their specialized formats for reference panel data: bref3 for BEAGLE5.1, m3vcf 4 for MINIMAC4 and imp5 for IMPUTE5. IMPUTE5 has two different haplotype selection algorithms that we call *divergence selection* and *neighbour selection* (see Methods), both of which have a parameter *L* that controls the number of selected haplotypes. We tested both selection algorithms using *L* = 4 and *L* = 8. IMPUTE4 was run with all reference panels except on the simulated reference panels, beceause IMPUTE4 is limited to 65,536 reference haplotypes and does not run on the two largest reference panels (100K and 1M samples).

As in previous papers [10, 12, 19], we measured performance by comparing the imputed allele probabilities to the true (masked) alleles. Markers were binned into bins according to the minor allele frequency of the marker in the reference panel. In each bin we report the squared correlation (*r*^2^) between the vector of all the true (masked) alleles and the vector of all posterior imputed allele probabilities.

All imputation analyses were run on a 16-core computer with Intel Xeon CPU E5-2667 3.20GHz processors and 512 GB of memory.

#### Imputation accuracy

Fig 2 shows the performance of all the methods on the Panel A simulated reference panels of size 10K, 100K, and 1M samples using *L* = 4 and *I* = 0.002cM for IMPUTE5. Results using *L* = 8 are shown in Figure S1. These results illustrate the very close agreement between the methods. All the methods compared use the same Li and Stephens probabilistic model [22] so this is not surprising. The imputation performance increases as expected with the reference panel size. For example, the imputation accuracy of the 10K reference panel at the 10^−4^ MAF bin reaches a *r*^2^ ≈ 0.4, while reaches a *r*^2^ ≈ 0.9 and *r*^2^ ≈ 0.98 for the 100K and 1M reference panel, respectively.

**Fig 2.**
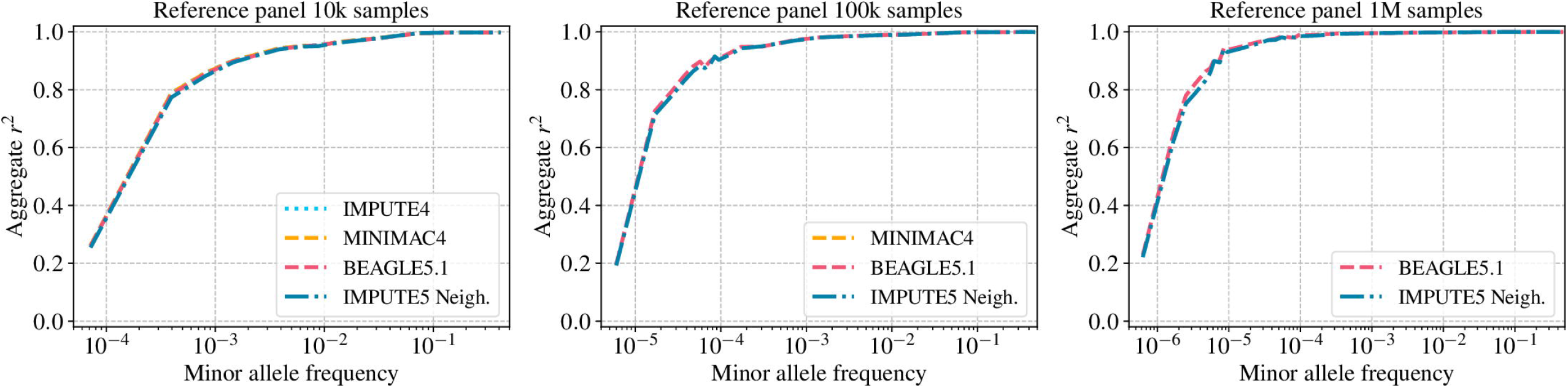
Imputation accuracy for the Panel A dataset. Imputation accuracy when imputing genotypes from a simulated reference panel of 10K, 100K and 1M UK-European reference samples (Panel A dataset). Imputed alleles are binned according to their minor allele frequency in each reference panel. The horizontal axis in each panel is on a log scale.

Fig 3 shows the results using the real reference panels and shows a very slightly increase in accuracy when MINIMAC4 is used for small reference panels. The likely explanation is that MINIMAC4 performs an HMM parameter estimation step when m3vcf files are created, and this adds some adaption to genotyping errors and recombination rate variation. This explains also why we do not see differences between methods in Fig 2, because in that case the real recombination map in known and no genotyping errors are present for the MSPRIME simulations. We also note that IMPUTE4 reaches the same imputation accuracy as other methods, even if run on smaller imputation windows. Results using *L* = 8 are shown in Figure S2.

**Fig 3.**
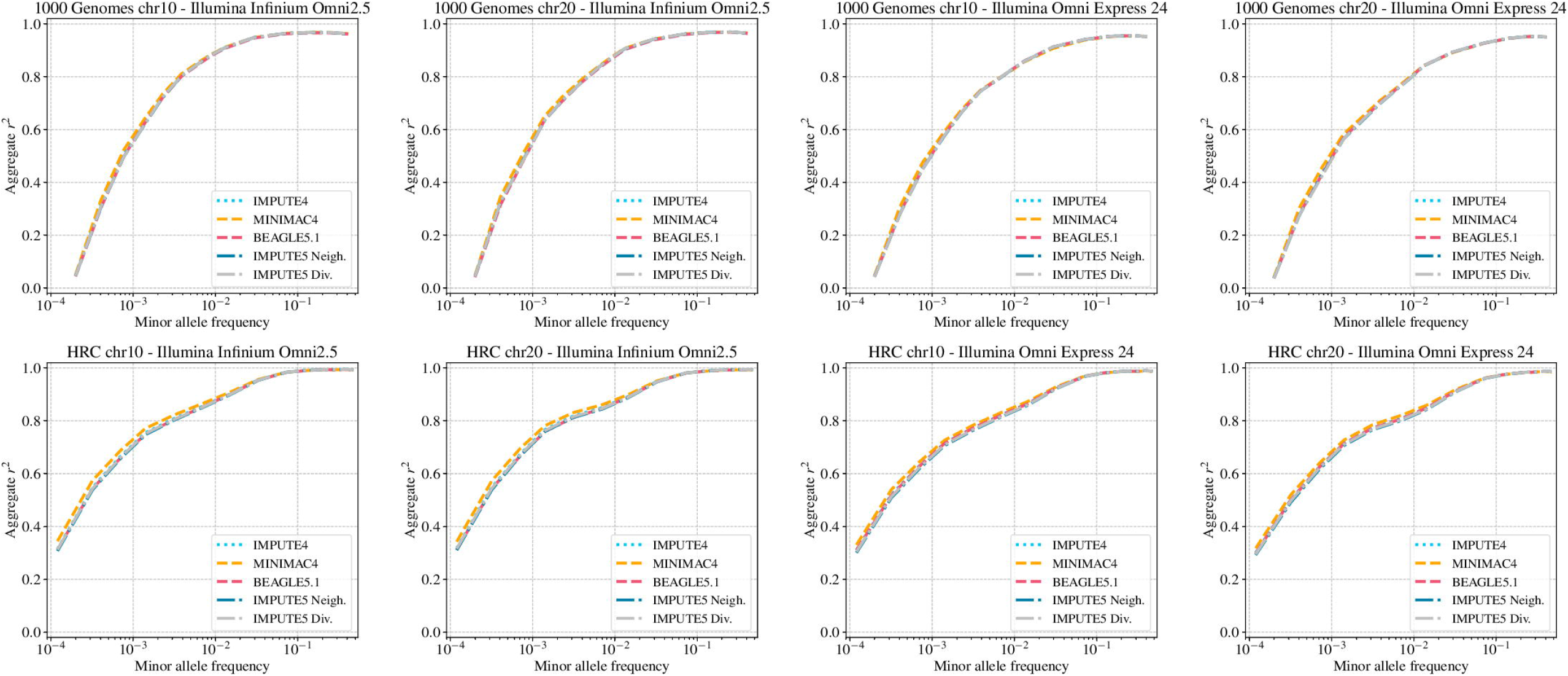
Imputation accuracy for the 1000 Genomes and the HRC datasets. Imputation accuracy when imputing genotypes using the 1000 Genomes Project reference panel (*n* = 2452) and the Haplotype Reference Consortium reference panel (*n* = 31470). Imputed alleles are binned according to their minor allele frequency in each reference panel. The horizontal axis in each panel is on a log scale.

Figure S3 and Figure S4 show the performance of IMPUTE5 for a range of values of *L* ∈ {1, 2, 4, 8, 16}) for both the selection algorithms proposed on simulated and real reference panels. As expected, increasing the value of *L* also accuracy increases, however, for values of *L* >= 4 imputation accuracy is almost indistinguishable. Both the selection algorithms perform well for values of *L* ≥ 4, however neighbour selection algorithm seems to perform better for values of *L* < 4. This is probably explained by the fact that neighbour selection algorithm tends to select more states than divergence selection algorithm, making it more robust even with smaller values of *L*.

We also verified the imputation accuracy in the case the target panel presents phasing errors. For this purpose we used our simulated datasets of Panel A., containing perfectly phased data. We added ≈ 2% switch error rate to each of the target datasets containing 1,000 samples. Overall, we note that a modest amount of phasing errors result in a drop in the imputation accuracy, especially in the rare frequency spectrum (Figure S5).

Finally, we used chromosome 10 data from the 1000 Genomes Project and HRC reference panel to explore the distribution of the selected states in our imputation experiments. We extracted 52 target samples from 1000 Genomes Project and 1000 target samples from the HRC, and phased them against the remaining haplotypes in the reference panels. We then recorded which haplotypes were selected as states across the ten 20cM chunks on chromosome 10. Figure S6 top shows the number of times each reference haplotypes was selected. The uneven pattern across reference haplotypes is a consequence of the spectrum of ancestry and different cohorts included in the 1000 Genomes and HRC reference panels respectively. Figure S6 bottom shows the distribution of the number of times a state was selected.

#### Computational efficiency

Table 4 shows single core memory usage and time of running MINIMAC4, BEAGLE5.1 and IMPUTE5 to impute the whole chromosome 20 and 10 for 1000 Genomes and HRC reference panels. In order to compare the methods, we run MINIMAC4 using its default chunk size (20 Mb chunk size and 2 Mb buffer size). IMPUTE5 and BEAGLE5 were run using 20 cM imputation regions and 1Mb and 2cM buffer respectively. IMPUTE5 was run with the parameter *L* = 4 and using BGEN as output file format (zstd compression). We used default settings otherwise. For both the 1000 Genomes and HRC reference panel IMPUTE5 is faster than BEAGLE5 and on the HRC reference panel it is over 20 times faster than MINIMAC4. IMPUTE5 is also several times more memory efficient than other methods. Results using *L* = 8 are shown in Table S1.

**Table 4.** Memory usage and time to impute 1000 Genomes and HRC datasets using 20Mb or 20cM imputation regions. Memory usage and total time to impute a whole chromosome (chr 10 and chr 20) for 52 target samples when using the 1000 Genomes reference panel and 1000 target samples when using the HRC reference panel. MINIMAC4 was run on chunks of size 20 Mb (default settings). IMPUTE5 and BEAGLE5.1 were run on chunks of size 20cM. Time is shown using the format mm:ss. Bold font is used to indicate the method with the lowest memory and time.

| Dataset | Memory usage (MB) - Omni 2.5 chip |  |  |  |
| --- | --- | --- | --- | --- |
|  | MINIMAC4 | BEAGLE5.1 | IMPUTE5 neigh | IMPUTE5 div |
| 1000 Genomes chr 20 | 1,792 | 6,706 | **428** | 435 |
| 1000 Genomes chr 10 | 2,472 | 12,089 | **480** | 489 |
| HRC chr 20 | 4,842 | 13,087 | **4,361** | 4,368 |
| HRC chr 10 | 4,896 | 14,480 | **4,842** | 4,850 |
|  | Memory usage (MB) - OmniExpress-24 chip |  |  |  |
|  | MINIMAC4 | BEAGLE5.1 | IMPUTE5 neigh | IMPUTE5 div |
| 1000 Genomes chr 20 | 1,696 | 6,232 | **197** | 200 |
| 1000 Genomes chr 10 | 2,316 | 11,572 | **215** | 221 |
| HRC chr 20 | 4,405 | 13,381 | **1,965** | 1,969 |
| HRC chr 10 | 4,417 | 13,351 | **1,779** | 1,788 |
|  | Time ([hh:]mm:ss) - Omni 2.5 chip |  |  |  |
|  | MINIMAC4 | BEAGLE5.1 | IMPUTE5 neigh | IMPUTE5 div |
| 1000 Genomes chr 20 | 02:33 | 00:55 | **00:48** | 00:49 |
| 1000 Genomes chr 10 | 05:16 | 01:44 | **01:42** | 01:42 |
| HRC chr 20 | 04:07:39 | 15:47 | **08:15** | 8:29 |
| HRC chr 10 | 08:07:42 | 30:47 | **17:27** | 17:52 |
|  | Time ([hh:]mm:ss) - OmniExpress-24 chip |  |  |  |
|  | MINIMAC4 | BEAGLE5.1 | IMPUTE5 neigh | IMPUTE5 div |
| 1000 Genomes chr 20 | 01:37 | 00:46 | **00:28** | 00:28 |
| 1000 Genomes chr 10 | 01:56 | 01:33 | **00:59** | 01:00 |
| HRC chr 20 | 02:02:29 | 11:03 | **03:38** | 03:43 |
| HRC chr 10 | 04:04:04 | 20:28 | **07:52** | 08:02 |

Fig 4 and Table 5 show the per-sample computation times for IMPUTE5, BEAGLE5.1 and MINIMAC4 for 10K, 100K and 1M simulated reference panels when imputing a set of 1,000 target samples on a 10Mb region. All methods were run using a single core on the same machine and in this case we reduced the value of the IMPUTE5 *I* parameter to 0.002 cM to take into account the fact that no proper map is available for the simulated region. The results are plotted on log-log scale, which illustrates that both BEAGLE5.1 and IMPUTE5 exhibit sub-linear scaling as reference panel size increases. For Panel A results, moving from 10K to 1M reference samples increases the number of reference samples by a factor of 100 and the number of reference markers by a factor of 10, but IMPUTE5’s imputation time increases by only a factor of 2.5. Overall the results show that IMPUTE5 is consistently faster than all the alternative methods. Results using *L* = 8 are shown in Figure S7 and Table S2.

**Fig 4.**
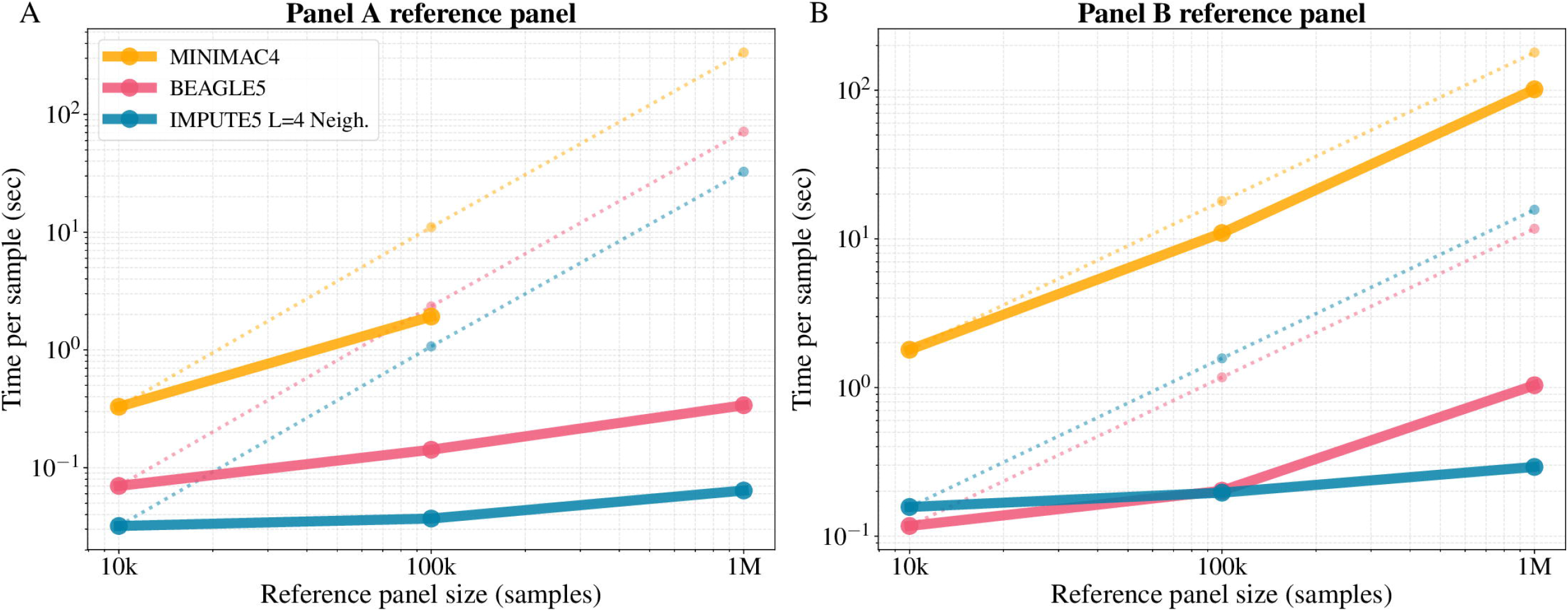
Per sample imputation time for Panel A and Panel B datasets. Per-sample CPU time when imputing a 10 Mb region from 10K, 100K and 1M simulated UK-European reference samples into 1,000 target samples using one computational thread per job. (A) Imputation time when using Panel A dataset (3,333 target markers). (B) Imputation time when using Panel B dataset (33,333 target markers). Axes are on log scale. Hypothetical linear scaling of MINIMAC4, BEAGLE5 and IMPUTE5 are shown as dotted lines, generated by projecting the time using the 10K reference panel. Minimac4 was not able to run using the Panel A 1M reference panel due to time constraints in the construction of the m3vcf file.

**Table 5.** Memory usage and time to impute Panel A and Panel B datasets. Memory usage and total time to impute 1, 000 target samples in a 10 Mb window using simulation data in Panel A and Panel B dataset. Time is shown using the format [hh:]mm:ss. Bold font is used to indicate the method with the lowest memory and time. MINIMAC4 was not able to run using the Panel A 1M samples reference panel due to time constraints in the construction of the m3vcf file.

| Dataset | Memory usage (MB) |  |  |  |  |  |
| --- | --- | --- | --- | --- | --- | --- |
|  | Panel A |  |  | Panel B |  |  |
|  | MINIMAC4 | BEAGLE5.1 | IMPUTE5 | MINIMAC4 | BEAGLE5.1 | IMPUTE5 |
| 10K reference panel | 1,048 | 9,058 | **1,210** | **1,253** | 7,278 | 12,241 |
| 100K reference panel | 5,858 | 10,452 | **1,283** | **5,122** | 7,994 | 11,842 |
| 1M reference panel | - | 8.805 | **2,217** | 43,246 | 25,868 | **15,659** |
| Dataset | Time ([hh:]mm:ss) |  |  |  |  |  |
|  | Panel A |  |  | Panel B |  |  |
|  | MINIMAC4 | BEAGLE5.1 | IMPUTE5 | MINIMAC4 | BEAGLE5.1 | IMPUTE5 |
| 10K reference panel | 05:30 | 01:11 | **00:32** | 29:54 | **01:58** | 02:39 |
| 100K reference panel | 32:20 | 02:23 | **00:36** | 03:02:21 | 03:23 | **03:20** |
| 1M reference panel | - | 05:39 | **01:17** | 28:10:50 | 17:15 | **05:51** |

Fig 4B and Table 5 show the imputation time for Panel B dataset. All the reference panels in Panel B have the same number of markers. In this case we have a very dense set target markers (33,333) and so more time is spent for the Li and Stephens calculations compared to Panel A scenario. The time spent by IMPUTE5 for the Li and Stephens HMM and imputation actually decreases when the number of reference haplotypes is increased. The increase in time shown in Fig 4B from 10K reference panel to 1M reference panel is only due to increased time to read the input and run the selection algorithm, the only linear components of IMPUTE5.

Fig 5A shows that the imputation time per sample decreases when the number of target haplotypes increases This is mainly explained by the fact that typically, for a small number of target haplotypes, the pbwt construction is the main part of the selection algorithm and the copying states selection is a small fraction of the time. Results using *L* = 8 are shown in Figure S8A.

**Fig 5.**
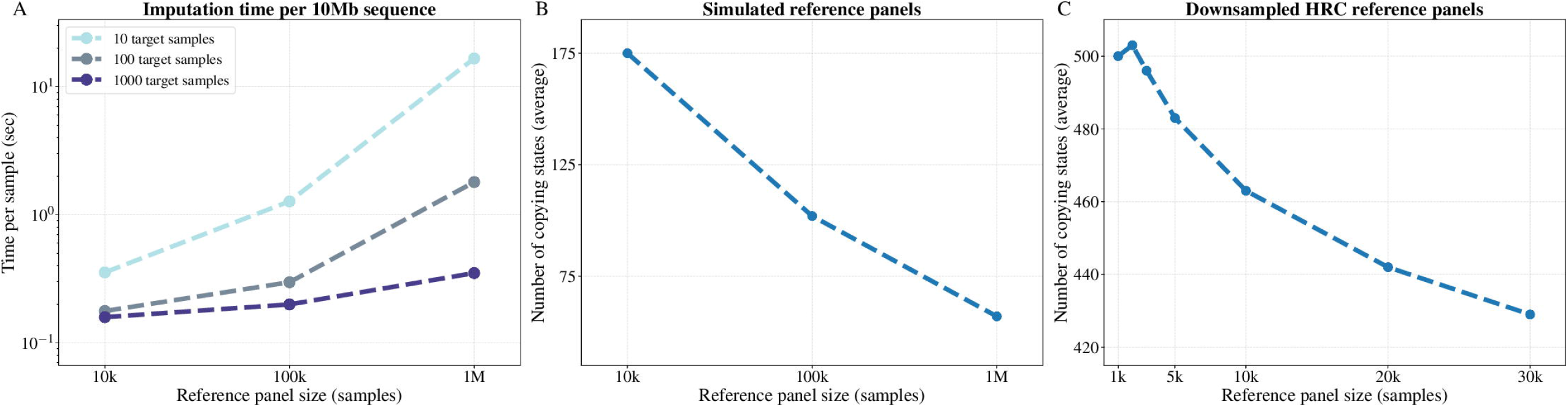
Sub-linear scaling of IMPUTE5. (A) Time per sample spent to impute a marker in a 10Mb region for reference panel size 10K, 100K and 1M samples, when imputing 10, 100 and 1,000 target samples. The vertical axis is on a log scale. (B) Mean number of copying states selected for the simulated reference panels. (C) Mean number of copying states selected for the downsampled HRC reference panels containing 1,000 to 30,000 thousands samples. Time and number of conditioning states are obtained using IMPUTE5 neighbours select and *L* = 4.

Fig 5B and C show that the mean number of copying states selected by IMPUTE5 decreases as the number of reference haplotypes increases for both simulated and real reference panels. For real reference panels, we used a downsampled version of the HRC containing an increasing amount of samples from 1,000 to 30,000. This property is predicted from the Li and Stephens model Eq 9, that is itself an approximation to the coalescent model, whereby the probability of switching between copying states decreases as the number of reference haplotypes increases. Results using *L* = 8 are shown in Figure S8B.

IMPUTE5 takes advantage of the BGEN file format. This is especially useful for very dense reference panels, containing millions of markers in a single chunk. In this case, we observe a 20 to 50% additional increase of speed, compared to using VCF/BCF file formats.

## Discussion

In this work we have developed a new genotype imputation method called IMPUTE5 that has the same accuracy and faster computation time and memory requirements compared to other currently available imputation methods. IMPUTE5 has the lowest computation time for all reference panel sizes and target sample sizes considered, both for small regions and for chromosome wide imputation.

IMPUTE5 shares the same model as IMPUTE4, but has several improvements, making IMPUTE5 suitable for new generation reference panels. A new reference file format (imp5) and the ability to read indexed input files allows quick imputation on a small region of the genome. IMPUTE5’s new copy states selection makes imputation more efficient when increasing the reference panel or target panel size. IMPUTE5 exhibits sub-linear scaling with reference panel sample size and provides highly accurate imputation for large scale data sets.

The ability to impute quickly specific regions of the genome makes IMPUTE5 very suitable to be used as a part of an imputation server [18]. In addition, IMPUTE5 could be optimized to be used after the pre-phasing step. For example, using SHAPEIT4 [6] to pre-phase target haplotypes using a reference panel of haplotypes, it internally computes the PBWT of the reference panel at target markers to provide an accurate phase. Since the same data structure is used in a similar way by the two programs, IMPUTE5’s selection algorithm could run as a last step of phasing.

We also believe that there is space for further improvements. For example, imp5 file format only provide a basic representation of the haplotypes and additional information can be added (i.e. PBWT divergence arrays). The ideas presented in this paper could be applied in other research areas, such as imputation for low coverage sequences, since the use of PBWT-based methods can improve speed and accuracy of imputation when imputing from a reference panel.

It seems likely that genotype imputation will continue to be an important part of most genome-wide association studies, since genotyping microarrays are relatively cheaper than whole-genome sequencing and as reference panels continue to grow.

Researchers will increasingly be able to impute (and re-impute) a larger number of rare variants and they will be imputed to a higher quality.

The increased length of haplotype matching that occurs as reference panels grow in size (see Fig 5B and C) suggests that it could be interesting to investigate whether genotyping microarrays could reduce the number of variants they assay without losing accuracy.

## Supporting information

Algorithm 1

Algorithm 2

## Software

The IMPUTE5 software is available at https://jmarchini.org/impute5/

## Acknowledgments

This work was supported by the UK Engineering and Physical Sciences Research Council (EPSRC) grant EP/G03706X/1 for the University of Oxford Doctoral Training Centre. We thank Prof. Pier Palamara for useful discussions and support.

## Supporting information

**S1 Algorithm. Divergence selection algorithm.**

**S2 Algorithm. Neighbour selection algorithm.**

**Table S1 Memory usage and time to impute 1000 Genomes and HRC datasets using** *L* = 8. Memory usage and total time to impute a whole chromosome (chr 10 and chr 20) for 52 target samples when using the 1000 Genomes reference panel and 1,000 target samples when using the HRC reference panel. MINIMAC4 was run on chunks of size 20 Mb while BEAGLE5 and IMPUTE5 on chunks of size 20 cM. Time is shown using the format mm:ss. Bold font is used to indicate the method with the lowest time.

**Table S2 Single core time to impute Panel A and Panel B datasets using** *L* = 8.Total time to impute 1, 000 target samples in a 10Mb window using simulation data in Panel A and Panel B dataset. Time is shown using the format mm:ss. Bold font is used to indicate the method with the lowest time. Minimac4 was not able to run using the Panel A 1M reference panel due to time constraints in the construction of the m3vcf file.

**Figure S1 Imputation accuracy for the Panel A dataset and** *L* = 8. Imputation accuracy when imputing genotypes from a simulated reference panel of 10K, 100K and 1M UK-European reference samples (Panel A). The horizontal axis in each panel is on a log scale.

**Figure S2 Imputation accuracy for the 1000 Genomes and the HRC datasets using** *L* = 8.Genotype imputation accuracy when imputing genotypes using the 1000 Genomes Project reference panel (*n* = 2452) and the Haplotype Reference Consortium reference panel (*n* = 31470). The horizontal axis in each panel is on a log scale.

**Figure S3 Imputation accuracy varying parameter** *L* **and the selection algorithm using Panel A dataset.** Genotype imputation accuracy when imputing genotypes using the 1000 Genomes Project reference panel (*n* = 2452) and the Haplotype Reference Consortium reference panel (*n* = 31470) for diffent values of the parameter *L* using the neighbour selection algorithm and the divergence selection algorithm. The horizontal axis in each panel is on a log scale.

**Figure S4 Imputation accuracy varying parameter** *L* **and the selection algorithm using 1000 Genomes and HRC datasets.** Imputation accuracy when imputing genotypes using the 1000 Genomes Project reference panel (*n* = 2452) and the Haplotype Reference Consortium reference panel (*n* = 31470) for different values of the parameter *L* using the neighbour selection algorithm and the divergence selection algorithm. The horizontal axis in each panel is on a log scale.

**Figure S5 Imputation performance in the case of phasing errors.** Imputation accuracy when imputing 1000 target samples from a simulated reference panel of 10K, 100K and 1M UK-European samples (Panel A) with no phasing errors (blue) and with a ≈ 2% switch error rate (red). The horizontal axis in each panel is on a log scale.

**Figure S6 Distribution of the selected states on real datasets.** Count of the number each reference haplotypes selected along the ten imputation chunks of chromosome 10 for the 1000 Genomes Project and HRC reference panel (top). Histogram of the selected counts for the two datasets (bottom).

**Figure S7 Per sample imputation time for Panel A and Panel B datasets and** *L* = 8.Per-sample CPU time when imputing a 10 Mb region from 10K, 100K and 1M simulated UK-European reference samples into 1,000 target samples using one computational thread. (A) Imputation time when using Panel A dataset (3,333 target markers). (B) Imputation time when using Panel B dataset (33,333 target markers). Axes are on log scale. Hypothetical linear scaling of MINIMAC4, BEAGLE5 and IMPUTE5 are shown as dotted lines, generated by projecting the time using the 10K reference panel. Minimac4 was not able to run using the Panel A 1M reference panel due to time constraints in the construction of the m3vcf file.

**Figure S8 Sub-linear scaling using** *L* = 8.(A) Time per sample spent to impute a marker in a 10Mb region for reference panel size 10K, 100K, 1000K, when imputing 10, 100 and 1000 target samples. The vertical axis is on a log scale. (B) Mean number of copying states selected for the simulated reference panels. The number of selected states decreases by increasing the size of the reference panel, showing sub-linear scaling. Time and number of conditioning states are obtained with neighbours select and *L* = 8.

