## Supplementary figures and images for "Genotype imputation using the Positional Burrows Wheeler Transform"

### Algorithm 1

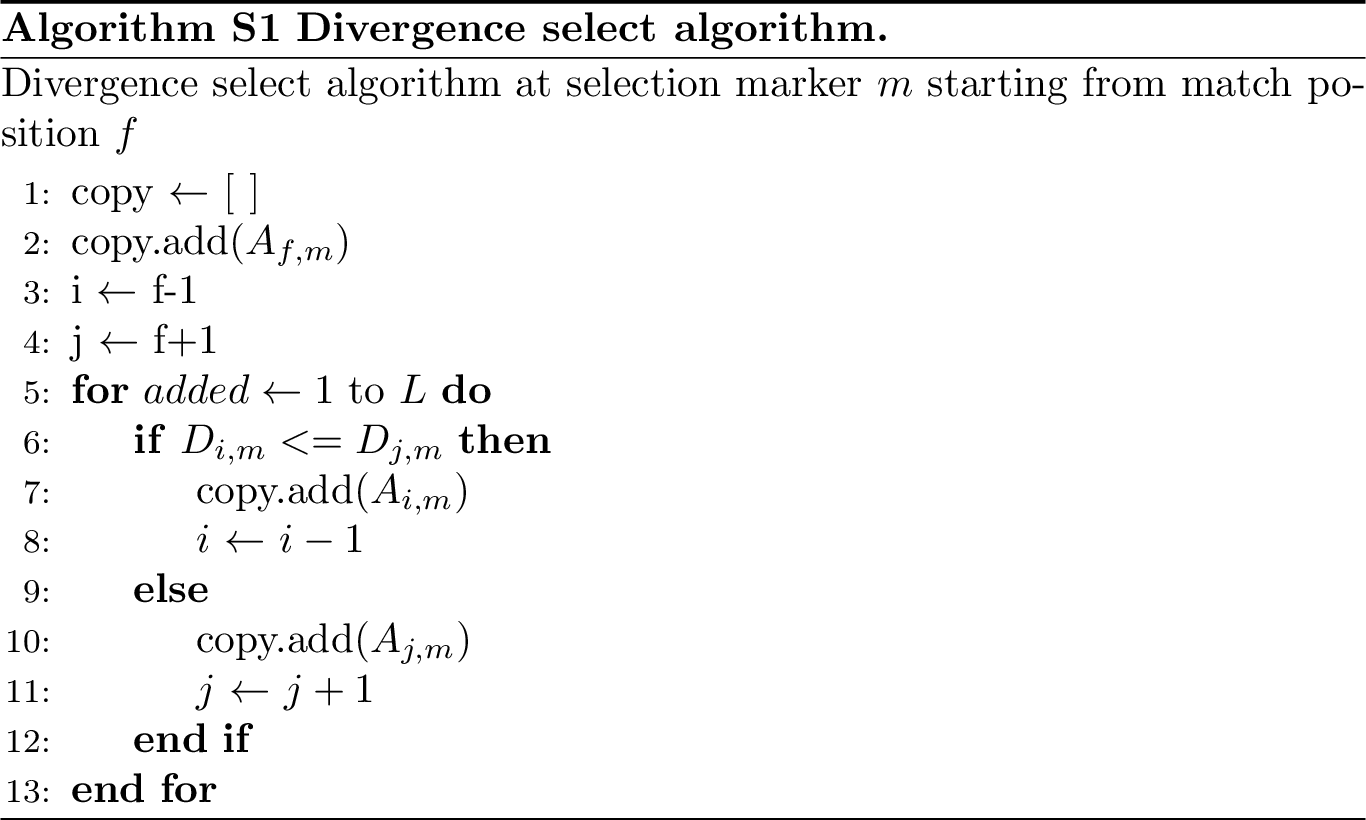

### Algorithm 2

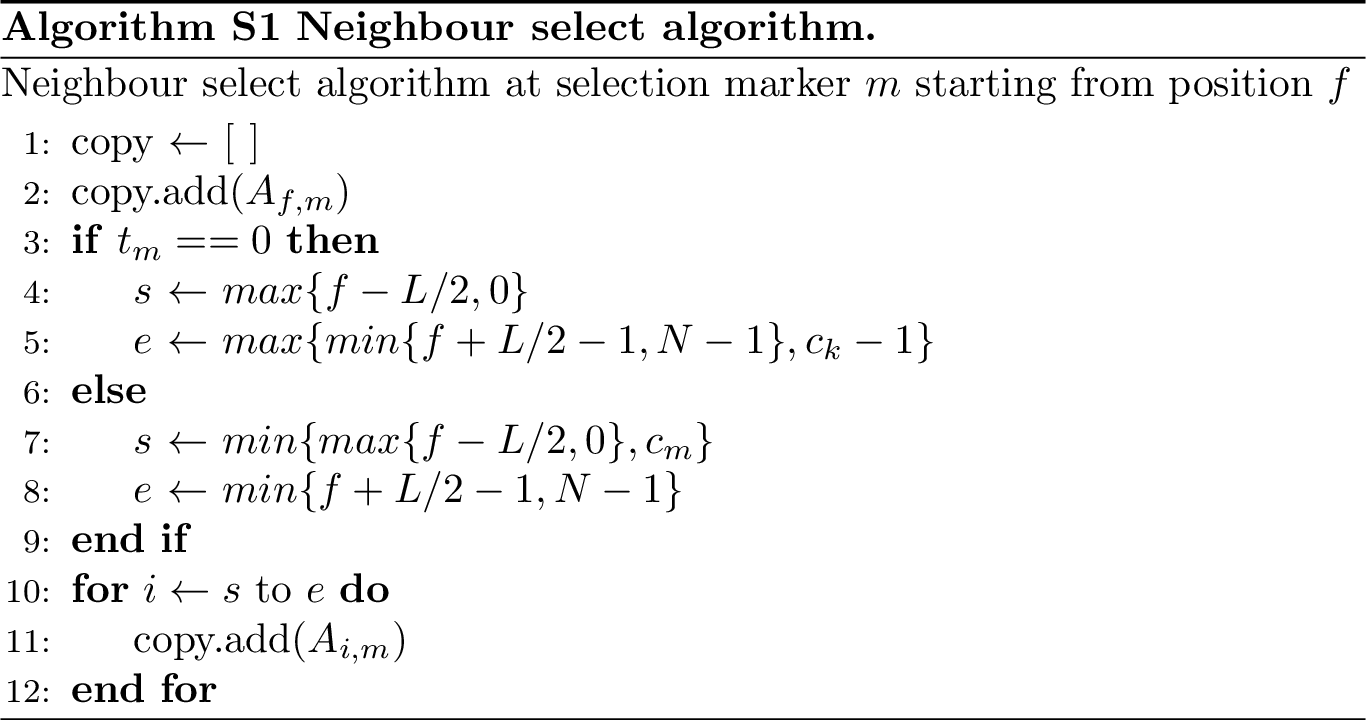

### Supplementary Figure 1

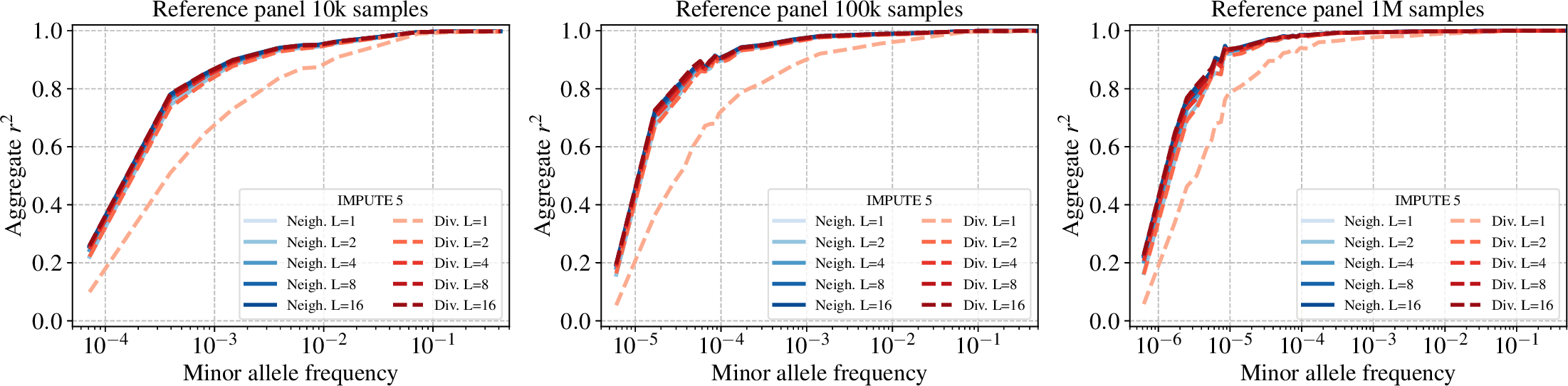

### Supplementary Figure 2

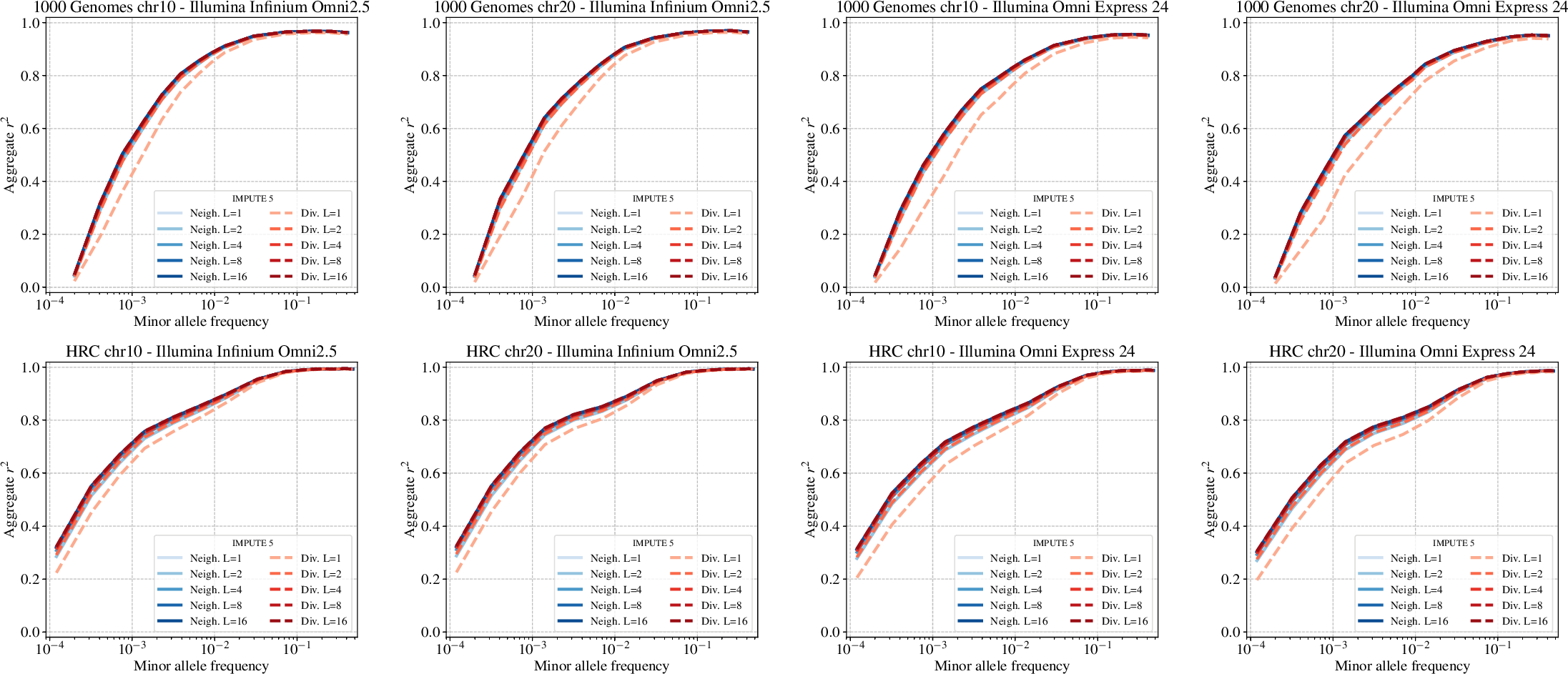

### Supplementary Figure 3

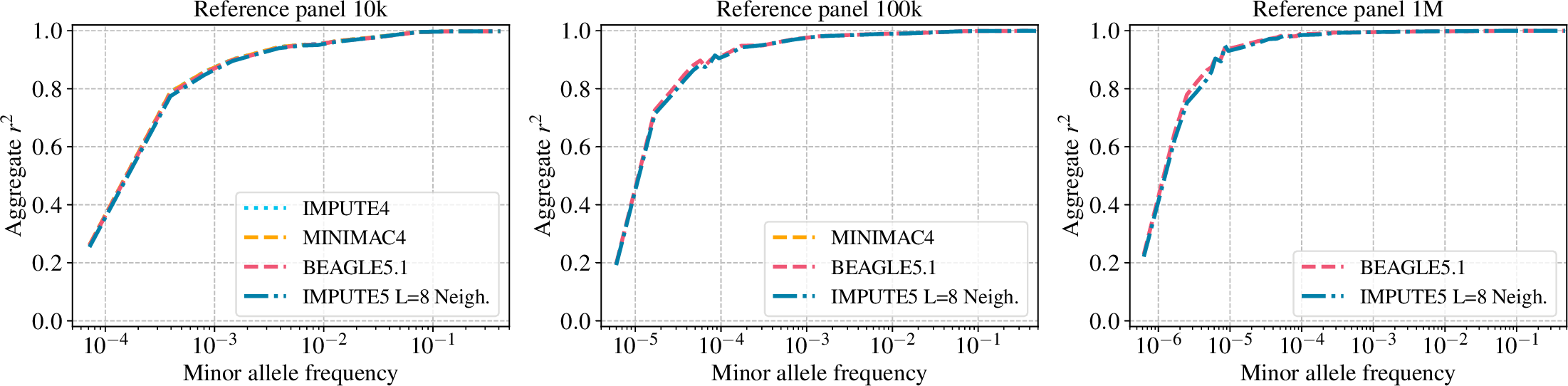

### Supplementary Figure 4

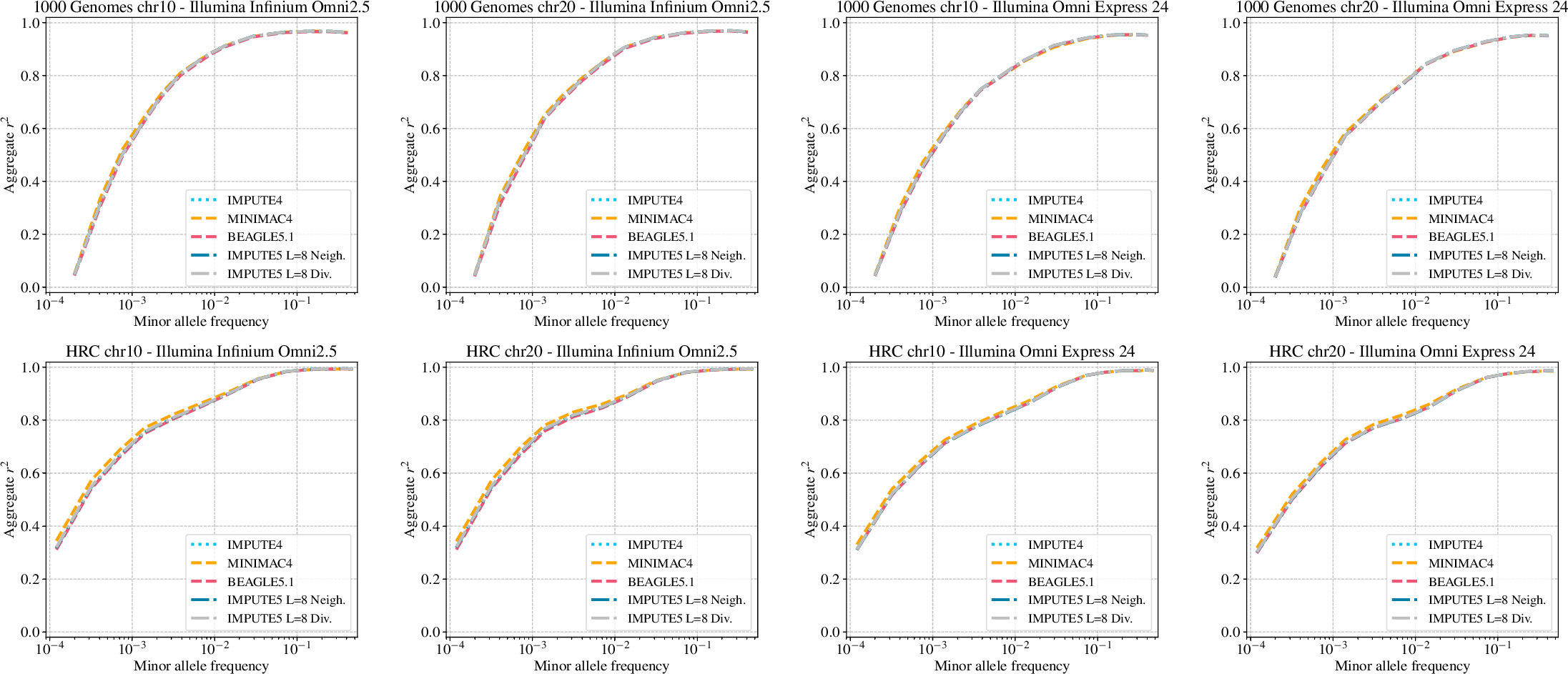

### Supplementary Figure 5

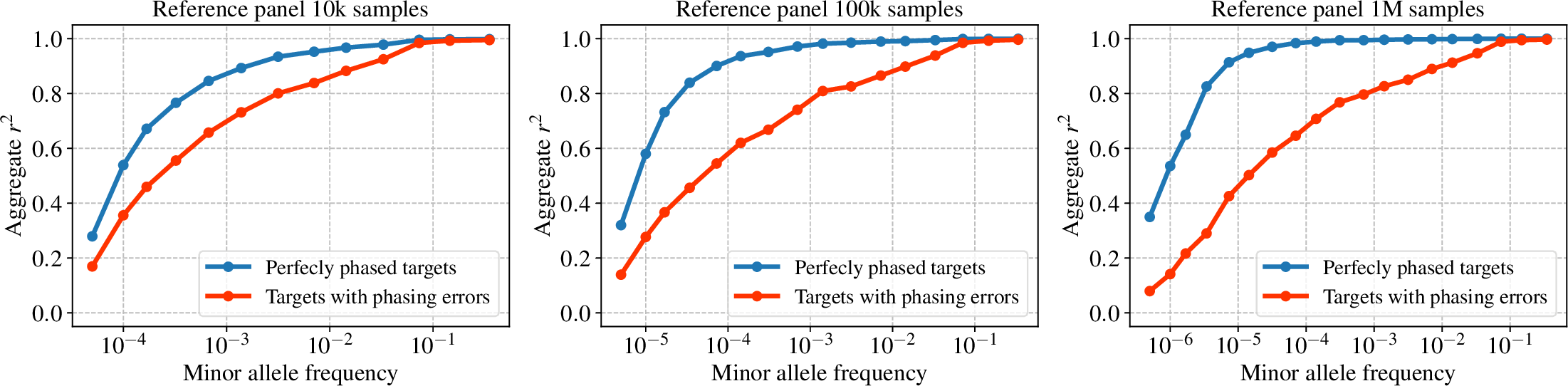

### Supplementary Figure 6

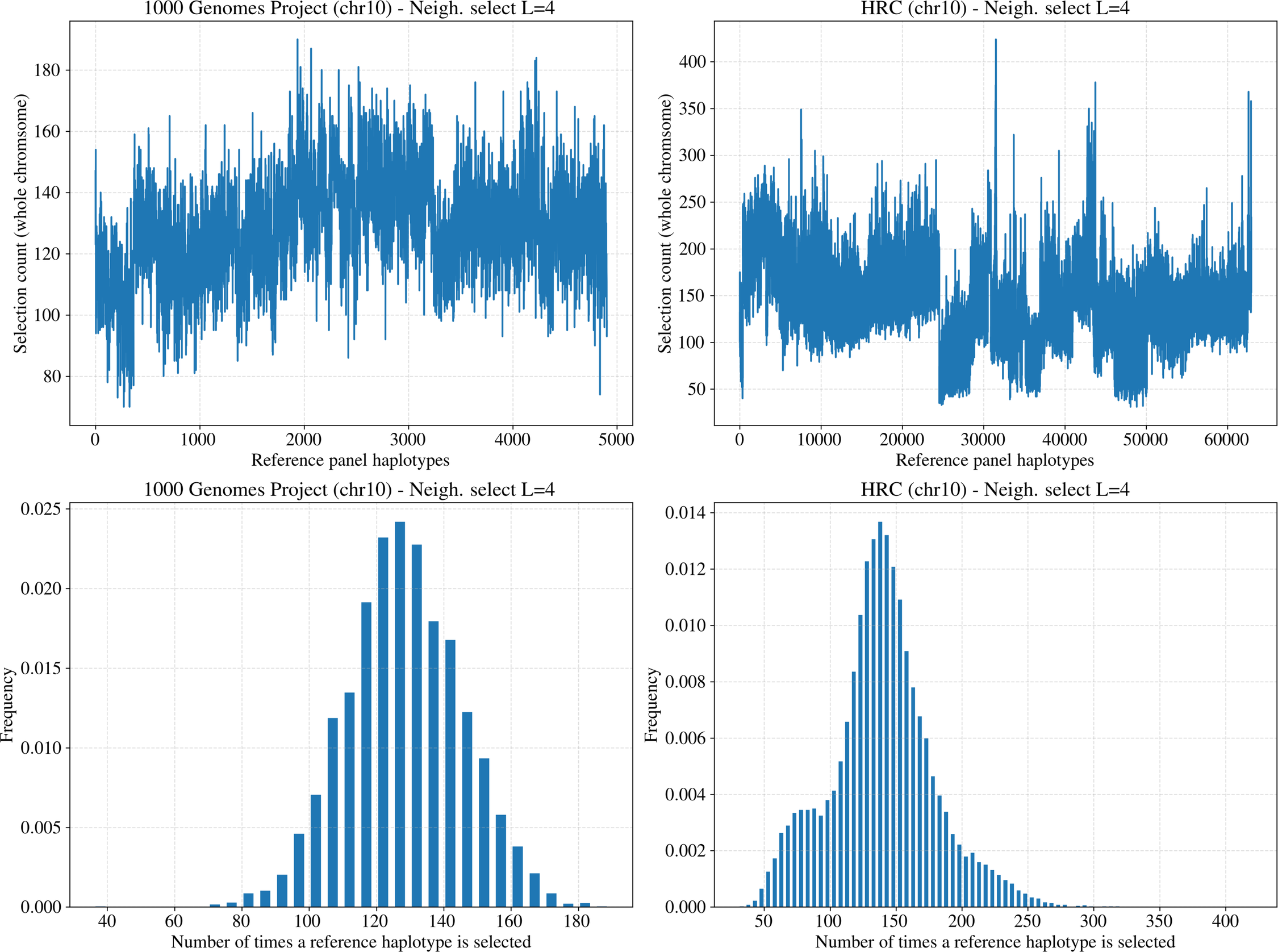

### Supplementary Figure 7

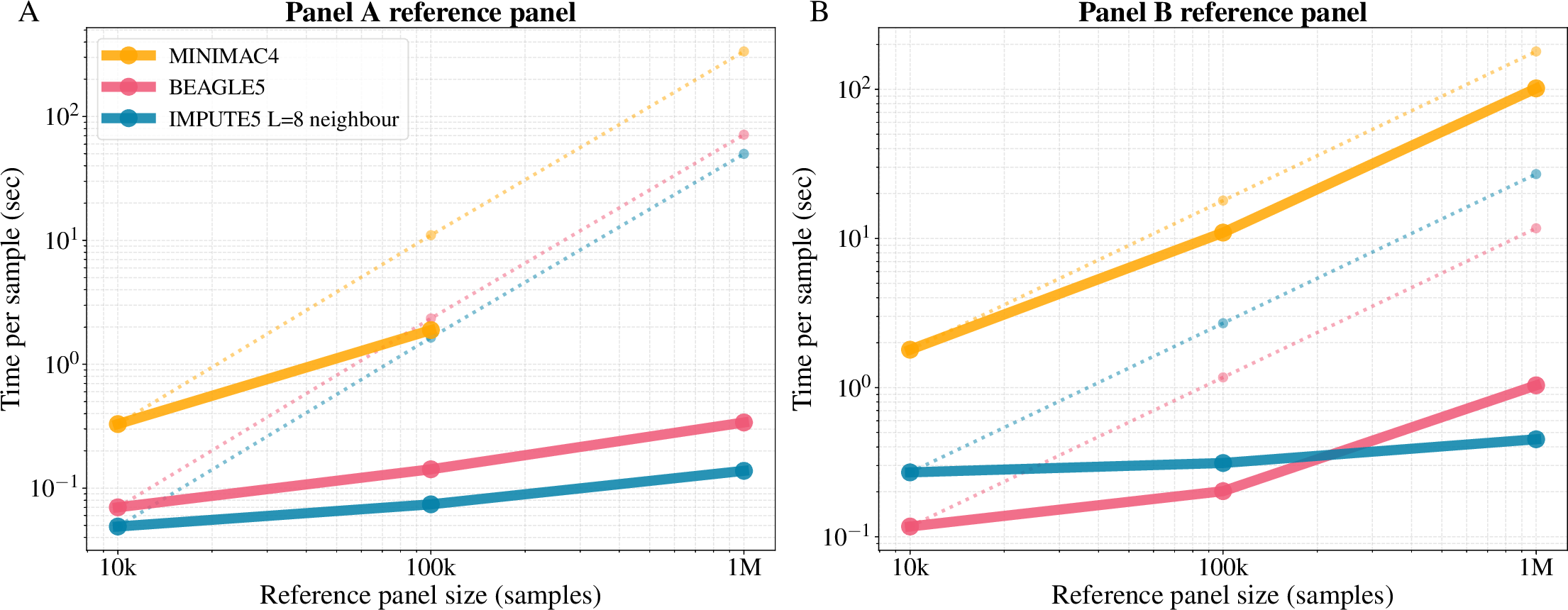

### Supplementary Figure 8

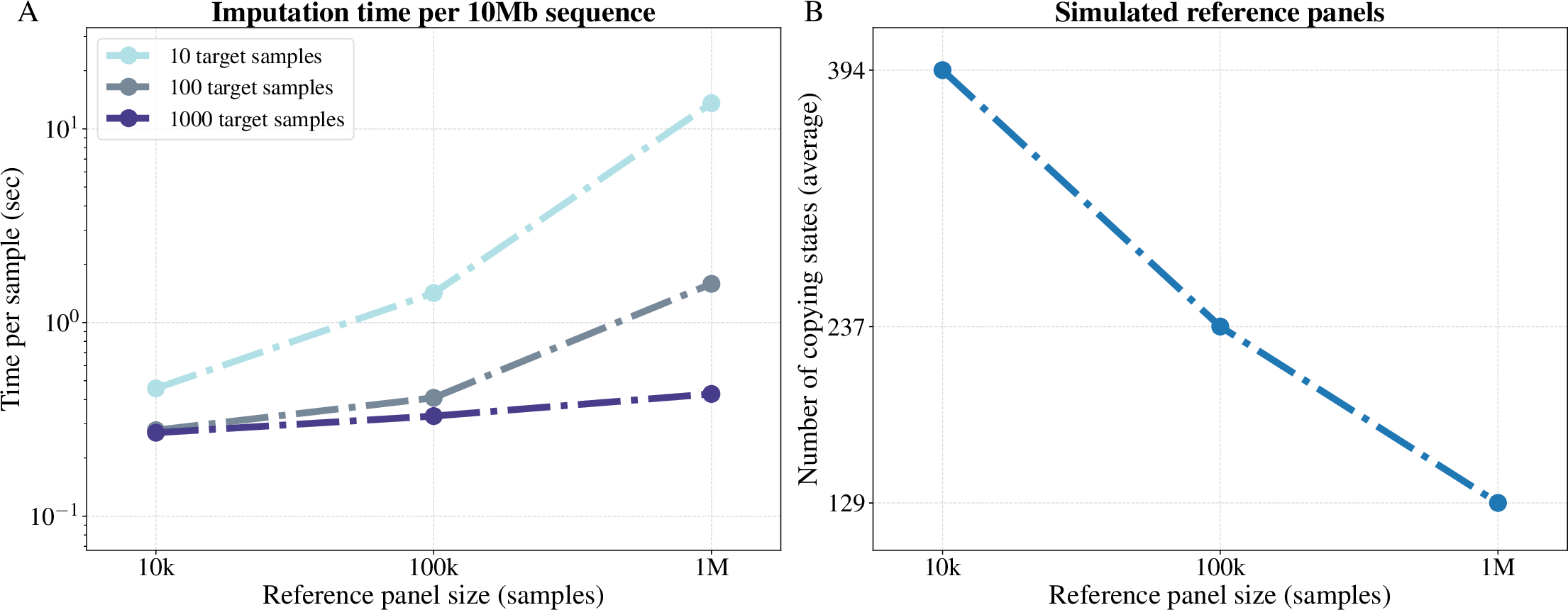

### Supplementary Table 1

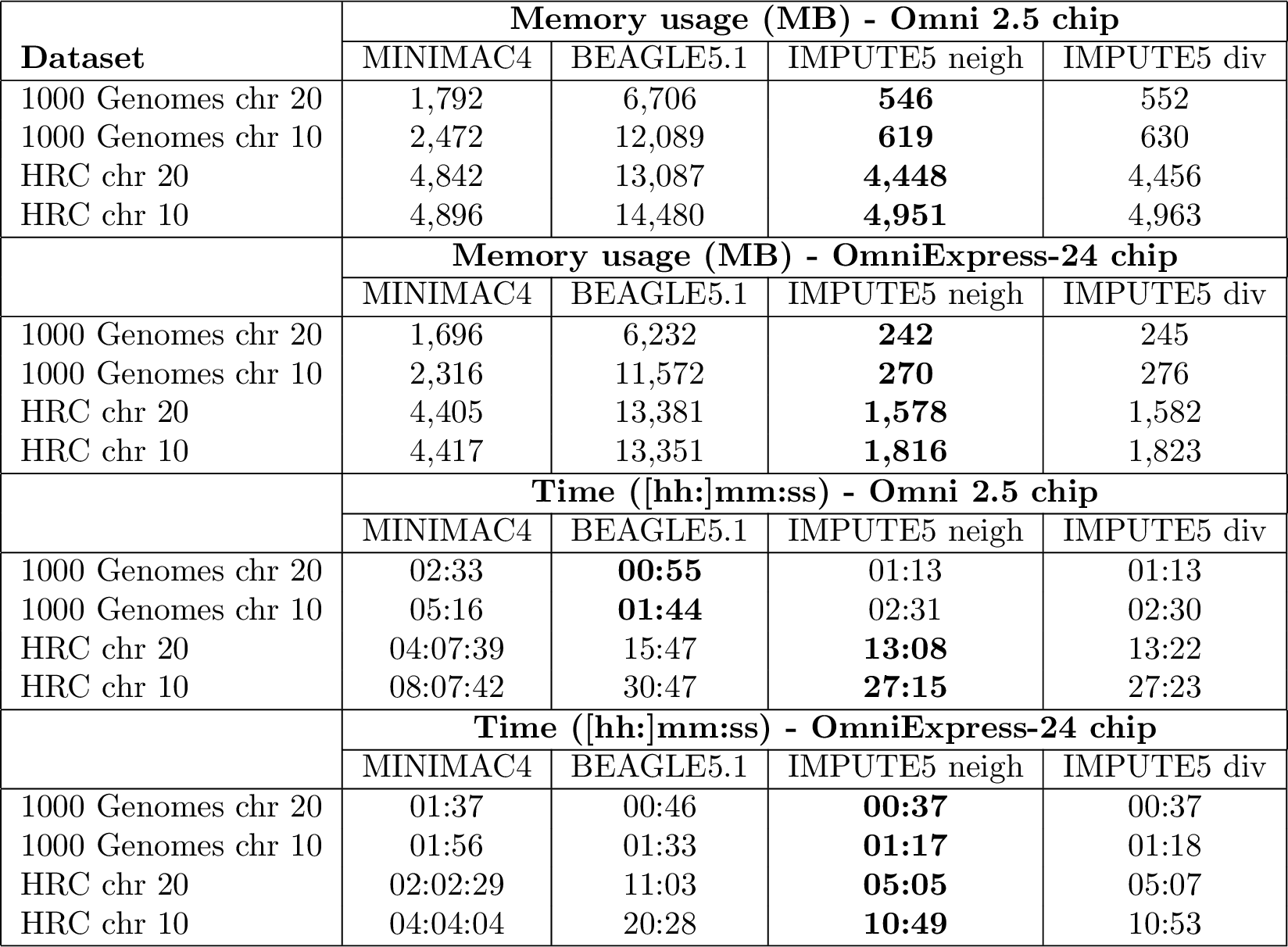

### Supplementary Table 2

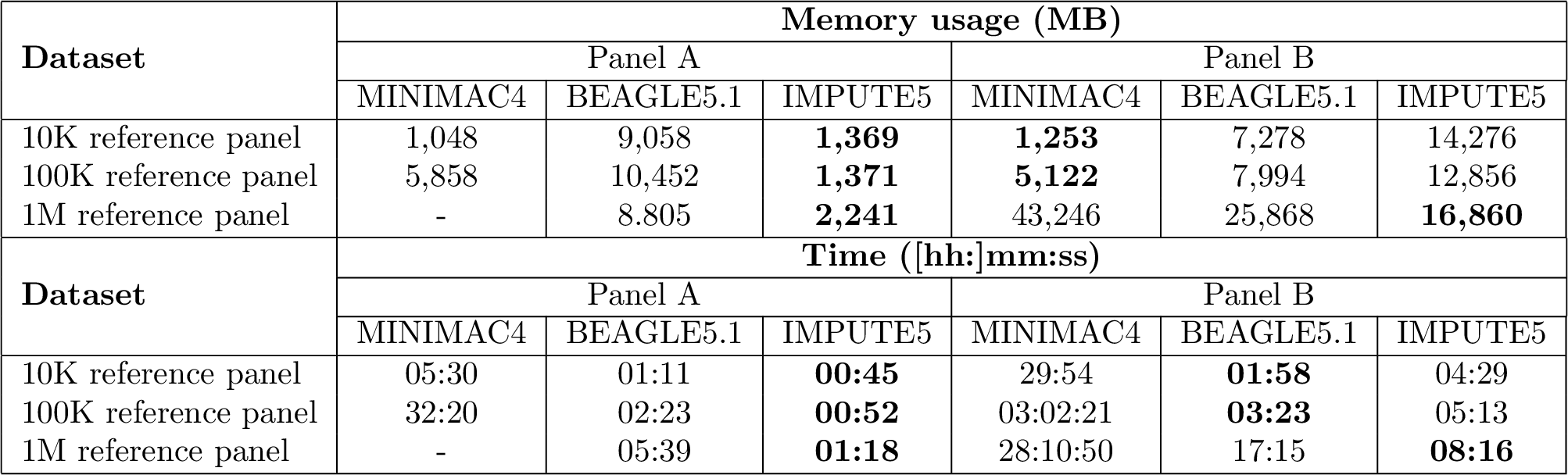
